# Magnetic Fields Improves the Cellular function in Hyperglycaemia

**DOI:** 10.64898/2026.08.30.748059

**Authors:** Bodhidipra Mukherjee, Shilpa Chandra, Abdul Salam, Laxmidhar Behera, Chayan Kanti Nandi

## Abstract

Hyperglycaemia disrupts mitochondrial homeostasis, leading to oxidative stress, ATP depletion, lysosomal dysfunction, impaired stress signalling, and proteostasis collapse. Although static magnetic fields (SMFs) have shown therapeutic potential in diabetic models, the optimal field strength for restoring subcellular organelle integrity post high glucose associated dysfunction remains unknown. Here, using Caenorhabditis elegans exposed to 40 mM glucose, we systematically evaluated SMFs ranging from 20 to 100 mT. Hyperglycaemia induced mitochondrial fragmentation, elevated reactive oxygen species, lysosomal abnormalities, reduced ATP levels, caused developmental delay, suppression of cytoprotective stress reporters, and increased polyglutamine aggregation. Among all field strengths tested, 70 mT produced the most robust recovery, restoring mitochondrial network architecture, reducing oxidative stress, normalizing lysosomal morphology, recovering ATP homeostasis, improving developmental progression, enhancing stress-responsive signalling, reducing proteotoxic aggregates, and increasing mitochondrial resilience to secondary hypoxic stress. These findings identify an optimal therapeutic SMF window and provide the first whole-organism demonstration that appropriately tuned static magnetic fields restore mitochondrial homeostasis and reverse multiple downstream consequences of hyperglycaemic stress.

## Introduction

Hyperglycaemia is one of the most significant metabolic disorders of the modern era and represents a major global health burden. The prevalence of diabetes mellitus has increased dramatically over the past few decades, affecting more than 500 million people worldwide, with projections indicating continued growth in the coming years [1]. Chronic elevation of blood glucose levels is associated with numerous complications including cardiovascular disease, nephropathy, retinopathy, impaired wound healing, accelerated ageing, and neurodegenerative disorders [2–6]. Although these pathologies manifest at the tissue and organismal level, their origins often arise from dysfunction at the subcellular level, particularly within mitochondria [7].

A variety of therapeutic approaches have been developed to mitigate hyperglycaemic damage. Sodium-glucose cotransporter-2 (SGLT2) inhibitors reduce blood glucose levels and provide cardiovascular benefits, while antioxidants limit oxidative stress and NAD+ boosters aim to enhance mitochondrial metabolism. However, prolonged use of SGLT2 inhibitors may increase the risk of mycotic and urinary tract infections [8], excessive NAD+ supplementation can cause digestive disturbances, allergies, and muscle fatigue [9], and overuse of antioxidants may interfere with chemotherapy efficacy in cancer patients [10]. Despite their promise, these interventions may exhibit off-target effects, variable efficacy, or limited long-term benefits. Therefore, alternative non-invasive strategies capable of restoring cellular energy homeostasis remain of considerable interest.

Static magnetic fields (SMFs) have emerged as a potential therapeutic modality capable of influencing biological systems without the need for pharmacological intervention [11,12]. Previous studies have demonstrated that magnetic fields can modulate mitochondrial activity, alter redox balance, influence ion transport, and affect cellular signalling pathways associated with metabolism and stress adaptation [13–16]. The radical pair mechanism further proposes that magnetic fields can influence ROS-generating biochemical reactions by altering the spin states of radical intermediates, thereby modulating oxidative stress responses [17]. Unlike conventional drug-based interventions, magnetic field therapy offers the possibility of restoring mitochondrial function and cellular physiology while minimizing systemic side effects. However, there is no general consensus regarding the optimal magnetic field strength or exposure duration required to reverse hyperglycaemia-induced organellar defects. A prominent study using diabetic mouse models reported that a 3 mT magnetic field combined with a 7 kV vertically oriented electric field reduced ROS levels and restored mitochondrial number. However, ROS decreased after 3 days of continuous exposure, whereas restoration of mitochondrial number required 30 days [18]. Another study used a 20–60 mT static magnetic field with continuous 24-hour exposure for 12–16 weeks in combination with intermittent fasting to reverse the autophagic burden associated with a high-fat diet in diabetic mice [19]. Similarly, continuous exposure to a 100 mT static magnetic field (24 hours/day) for 12 weeks restored blood glucose levels in diabetic mice [19].

However, these studies primarily focused on reducing ROS, restoring blood glucose, altering cytoprotective protein expression, or reversing body weight changes. Hyperglycaemia also causes extensive organellar damage, with mitochondria being one of its primary targets. Excess glucose increases electron flux through the mitochondrial electron transport chain, promoting electron leakage and excessive ROS generation [20,21]. Persistent oxidative stress damages mitochondrial proteins, lipids, and DNA, causing mitochondrial fragmentation, reduced ATP production, impaired stress signalling, disrupted proteostasis [22,23], lysosomal dysfunction [24], and accumulation of aggregation-prone proteins associated with Alzheimer’s, Parkinson’s, and Huntington’s diseases [25,26]. Consequently, restoration of mitochondrial function has emerged as an important therapeutic strategy.

The precise mechanism by which magnetic fields influence mitochondrial function remains unclear. However, several free radical-generating sites within the mitochondrial electron transport chain are thought to respond to magnetic field exposure via the radical pair mechanism [21]. Pulsed electromagnetic fields (PEMFs) have been reported to promote mitochondrial fission during angiogenesis in HUVEC cells, suggesting that electromagnetic stimulation can directly modulate mitochondrial dynamics. In contrast, static magnetic fields (SMFs) have been shown to reduce mitochondrial ROS under hyperglycaemic conditions [19,20]. Despite these encouraging findings, no standardized therapeutic protocol has been established regarding the optimal magnetic field strength or exposure duration required to achieve beneficial effects while avoiding potential off-target consequences at higher field strengths. Indeed, previous studies have employed a wide range of magnetic field intensities (20–100 mT) to lower blood glucose levels, enhance cytoprotective protein expression, reduce oxidative stress, and restore organ function in diabetic models, with treatment durations typically ranging from 4 to 12 weeks. However, none of these studies investigated the restoration of subcellular organelle morphology at the whole-organism level. Therefore, in the present study, we investigated whether static magnetic fields could reverse hyperglycaemia-induced organellar dysfunction, with particular emphasis on mitochondrial dysregulation and its downstream cellular consequences. Using C. elegans as an in vivo model of hyperglycaemia, we systematically evaluated multiple static magnetic field strengths and identified 70 mT as the optimal field strength, producing the most robust recovery across mitochondrial morphology, lysosomal integrity, oxidative stress, ATP homeostasis, proteostasis, and developmental phenotypes. These findings demonstrate the therapeutic potential of appropriately tuned static magnetic fields for reversing hyperglycaemia-induced cellular dysfunction.

## Results

### 1. Effect of High Glucose on *C. elegans* Mitochondria

Given the central role of mitochondria in maintaining cellular energy homeostasis, we first evaluated whether static magnetic fields (SMFs) could influence mitochondrial health under basal conditions before investigating their therapeutic potential during hyperglycaemic stress. Control worms exposed to a 70 mT SMF exhibited a slight, non-significant increase in overall mitochondrial area **(Figure S1a-i–iii),** and a significant increase in number of junctions (**(Figure S2)**. As previously discussed, healthy mitochondria exhibit a meshwork morphology large number of junctions hence the data suggests that this field strength may possess intrinsic mitochondrial protective properties without adversely affecting healthy tissue. We then examined the effects of hyperglycaemia and static magnetic field exposure on mitochondrial morphology in vivo. Exposure to 40 mM glucose resulted in a marked reduction in mitochondrial meshwork integrity compared to control populations, as qualitatively observed in **(Figure 1a-i–ii).** Progressive restoration of the mitochondrial network was subsequently observed with increasing static magnetic field strength from 20–70 mT **(Figure 1a-iii–v),** with the most pronounced recovery occurring at 70 mT. However, this beneficial effect was lost at higher field strengths, with worms exposed to 100 mT displaying a reduction in mitochondrial network distribution and increased fragmentation **(Figure S3a-i–iv),.** Quantitative analysis supported these observations **(Figure 1b-c).** Treatment with a 70 mT magnetic field significantly increased the average mitochondrial area from 0.70 to 1.05 μm² (p = 0.008) and perimeter from 3.86 to 5.10 μm (p = 0.007) compared to hyperglycaemic worms **(Figure 1b-i–ii).** These values approached those observed in control animals, which exhibited an average mitochondrial area of 1.08 μm² and perimeter of 5.34 μm. Hyperglycaemia also increased mitochondrial circularity from approximately 0.59 to 0.70, indicating a transition from elongated interconnected networks to fragmented punctate structures **(Figure S4),** however, the effect was reversed after exposure to 70mT magnetic field. Interestingly, the average number of detectable mitochondria increased from 594 to 796 following 70 mT treatment **(Figure 1d),** suggesting improved preservation or recovery of the mitochondrial network.

**Figure 1:**
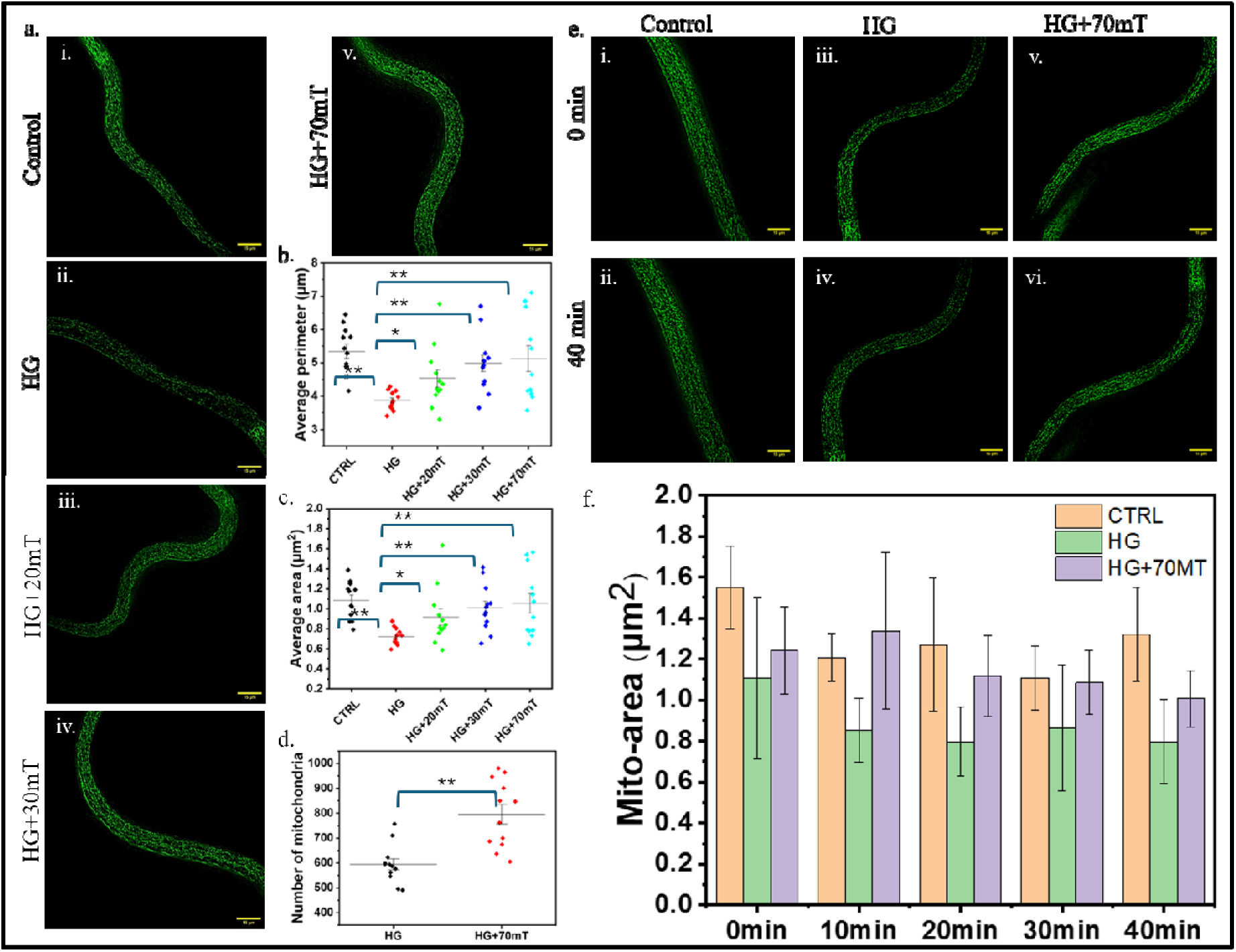
Effect of high glucose on C. elegans mitochondria and their regeneration via magnetic field. (a-i-v) Shows hyperglycaemia-induced disruption of the mitochondrial network in 40 mM glucose-exposed C. elegans and its progressive restoration with increasing magnetic field strength. (b,c) Quantitative comparison of mitochondrial area (b) perimeter (c) in hyperglycaemia-exposed C. elegans across increasing magnetic field strengths (d) Increase in mitochondrial number after exposure to magnetic field. (e, i-vi) Mitochondrial Area Changes During Hypoxia: Control vs High Glucose vs Magnetic Field Treatment. (f) Quantitative analysis of the time-dependant changes in area.

To further assess mitochondrial resilience under external stress, worms were immobilized in 50% (w/v) Pluronic F-127 polymer. At this concentration, the polymer creates a near-hypoxic environment while simultaneously facilitating long-term imaging. Time-course analysis of the same animals revealed a progressive decline in mitochondrial area over 40 minutes in all experimental groups, including control, hyperglycaemic, and magnetic-field-treated worms **(Figure 1e-i–vi).** However, hyperglycaemic animals exhibited substantially smaller mitochondria at the onset of the experiment and showed a more severe decline during hypoxic stress. The average mitochondrial area decreased from approximately 1.1 to 0.6 μm² over the 40-minute period, compared to final values of 1.3 μm² in control animals and 1.0 μm² in magnetic-field-treated hyperglycaemic animals **(Figure 1f).** These observations suggest that hyperglycaemia compromises mitochondrial resilience to secondary stressors, while magnetic field treatment partially restores mitochondrial stress tolerance. Together, these findings demonstrate that static magnetic fields promote restoration of mitochondrial architecture following hyperglycaemic damage while also enhancing mitochondrial network integrity under basal physiological conditions.

### 2. Effect of High Glucose on *C. elegans* ROS and lysosomal morphology

Given the close relationship between mitochondrial dysfunction and oxidative stress, we next investigated whether the mitochondrial alterations observed following hyperglycaemia were accompanied by changes in intracellular ROS levels. For this purpose, we employed the CYA19 strain expressing the gst-4p::GFP::NLS reporter. Glutathione S-transferase-4 (GST-4) is a key cytoprotective enzyme that is transcriptionally upregulated in response to elevated ROS levels [27]. Consistent with the mitochondrial dysfunction observed in hyperglycaemic worms, exposure to 40 mM glucose resulted in a significant increase in GFP fluorescence compared to control animals (p < 0.0001), indicating elevated oxidative stress throughout the body of C. elegans **(Figure 2a-i–ii).** Quantitative analysis further confirmed a substantial increase in reporter intensity following glucose exposure **(Figure 2b-i–ii).** Interestingly, treatment with a 70 mT static magnetic field significantly reduced gst-4 reporter fluorescence (p < 0.0001), restoring signal levels close to those observed in untreated controls **(Figure 2c-i–ii).** This observation is consistent with previous reports suggesting that magnetic fields can influence ROS-generating biochemical reactions through the radical pair mechanism while simultaneously promoting cytoprotective stress-response pathways [28]. Quantitative comparison revealed a marked reduction in overall reporter intensity following magnetic field treatment **(Figure 2d-i).** Similarly, the percentage of body area occupied by the reporter decreased from approximately 90% in hyperglycaemic worms to nearly 70% (p < 0.0001), following magnetic field exposure, approaching the control value of approximately 66% **(Figure 2d-ii).** Together, these findings indicate that static magnetic field treatment effectively mitigates hyperglycaemia-induced oxidative stress. Since persistent oxidative stress can impair organelle quality-control mechanisms, we next examined the effects of hyperglycaemia on lysosomal morphology and function. Hyperglycaemic worms exhibited pronounced lysosomal accumulation compared to control animals as seen in **(Figure 2e-i–iii),** where hyperglycaemic animals displayed substantially higher lysosomal burden than control worms. Notably, magnetic field treatment reduced lysosomal accumulation and restored a more dispersed lysosomal distribution pattern. Quantitative analysis revealed that the average lysosomal area increased significantly following glucose exposure from 0.41 to 0.48 μm², indicative of lysosomal enlargement and aggregation. Following magnetic field treatment, this value decreased to approximately 0.42 μm², closely resembling control levels **(Figure 2f-iv).** These observations suggest that restoration of mitochondrial homeostasis and reduction of oxidative stress by static magnetic fields are accompanied by a corresponding recovery of lysosomal integrity and function.

**Figure 2:**
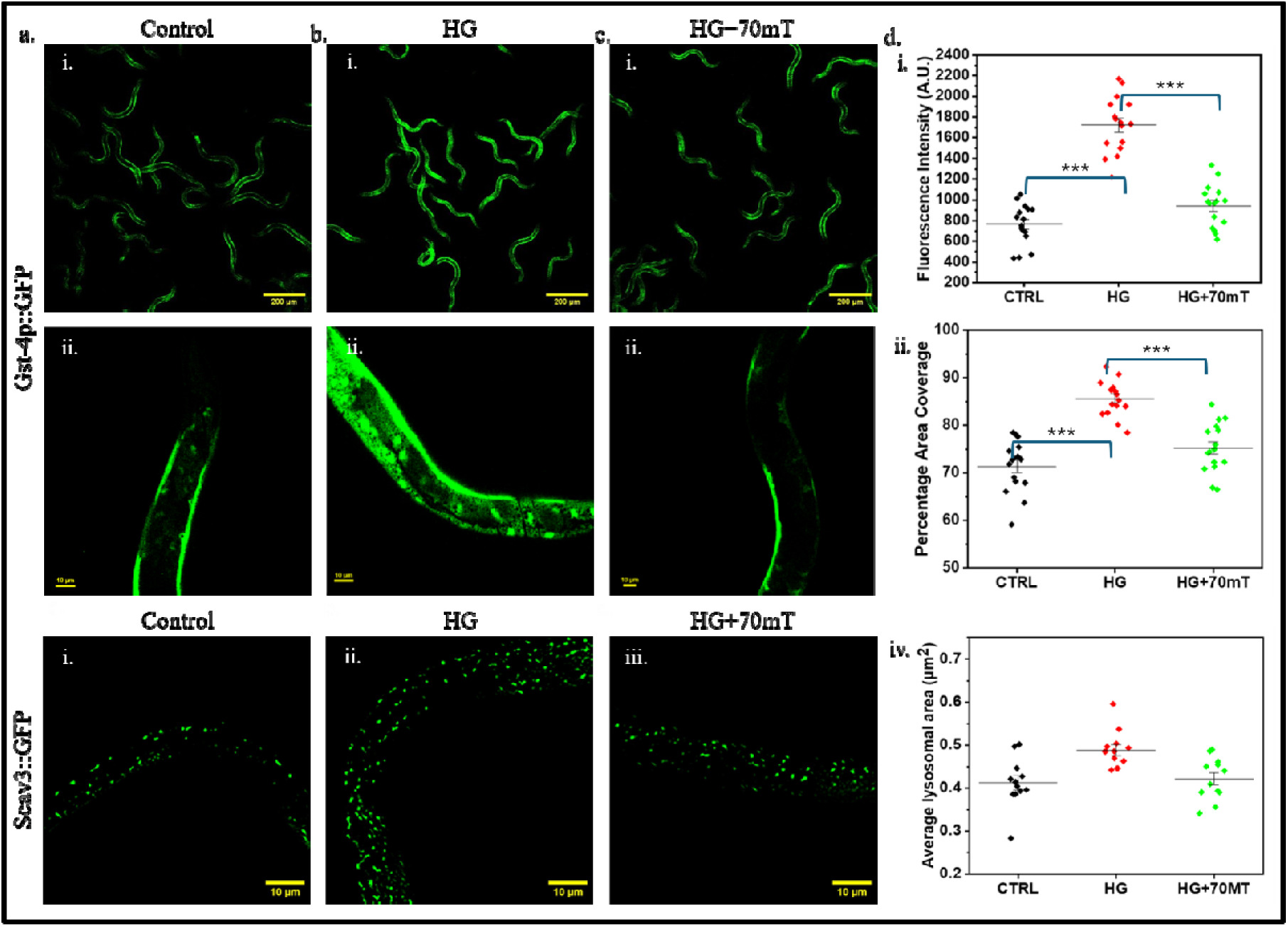
Effect of high glucose on C. elegans Ros sensor and lysosomal distribution. **(a)** Shows oxidative stress related Gst-4p fluorescence in the body of C. elegans in **(i)** overall population (**ii)** magnified version. **(b)** Shows oxidative stress related Gst-4p fluorescence in the body of C. elegans after exposure to high glucose in **(i)** overall population (**ii)** magnified version showing nuclear **(c)** Shows oxidative stress related Gst-4p fluorescence in the body of C. elegans in **(i)** overall population (**ii)** magnified version showing reduced nuclear localisation. **(d-i-ii)** Quantitative comparison of **(i)** fluorescence intensity **(ii)** percentage area coverage of Gst-4p based fluorescence within the body of C. elegans under different experimental conditions. **(f-i-iii)** Lysosomal distribution in C. elegans from control, high glucose exposed and magnetic field treated worms previously exposed to high glucose. **(d-iv)** Quantitative comparison of average lysosomal area in C. elegans from different experimental conditions.

### 3. Effect of mitochondrial dysfunction on cellular energy environment and its downstream consequence of C. elegans development

Given that lysosomal function, oxidative stress responses, and cellular maintenance are highly dependent on ATP availability, we next investigated whether the mitochondrial and lysosomal dysfunction observed under hyperglycaemic conditions translated into measurable changes in cellular energetics. For this purpose, we employed the NK3304 strain expressing the PercevalHR biosensor, which provides a real-time readout of cellular ATP ratios in living worms. The sensor functions through competitive binding of ATP and ADP to the GlnK1 nucleotide-binding domain fused to cpVenus, resulting in ratiometric fluorescence changes that reflect cellular energy status [29]. ATP binding increases fluorescence upon excitation at 500 nm, whereas ADP binding enhances fluorescence following excitation at 420 nm, allowing quantitative assessment of ATP ratios [29]. Consistent with the mitochondrial fragmentation observed in previous sections, hyperglycaemic worms exhibited a markedly reduced ATP ratio compared to control animals **(Figure 3a-i–ii, b-i–ii).** This reduction likely reflects impaired mitochondrial bioenergetics despite the abundance of extracellular glucose. Interestingly, magnetic field treatment restored cellular energy levels **(Figure 3c-i–ii),** in agreement with the improvements in mitochondrial morphology, oxidative stress, and lysosomal integrity described above. Quantitative analysis revealed that the average ATP ratio increased significantly from 3.3 in hyperglycaemic worms to 7.1 following 70 mT magnetic field treatment (p = 0.002), approaching the control value of 7.8 **(Figure 3d).** The response was also field-strength dependent, with 30 mT producing only partial recovery and an average ATP ratio of 5.3. Because cellular energy availability is essential for growth and development, we next examined whether these metabolic alterations affected organismal development. Wild-type N2 animals were imaged 48–54 hours following age synchronization, a developmental stage at which most animals are expected to have reached the late L4 or young adult stage. This developmental progression was observed in control populations; however, worms exposed to high glucose displayed a pronounced developmental delay and remained predominantly at the L3–L4 stages **(Figure 3e-i–iii).** Remarkably, exposure to a 70 mT magnetic field largely rescued this phenotype, resulting in developmental progression comparable to control animals. Population-level body size measurements further supported these observations. Control worms exhibited an average body length of approximately 834 μm, whereas hyperglycaemic animals were significantly smaller, with an average length of nearly 616 μm (p < 0.0001). Magnetic field treatment restored growth, increasing average body length to approximately 820 μm, closely matching control values **(Figure 3f).** Together, these findings indicate that hyperglycaemia-induced mitochondrial dysfunction results in a profound energetic deficit that impairs normal development, while static magnetic field treatment restores cellular bioenergetics and promotes recovery of organismal growth. Given the central role of ATP in maintaining protein quality-control pathways, we next investigated whether these alterations in cellular energy status affected other stress-response signalling and proteostasis pathways.

**Figure 3:**
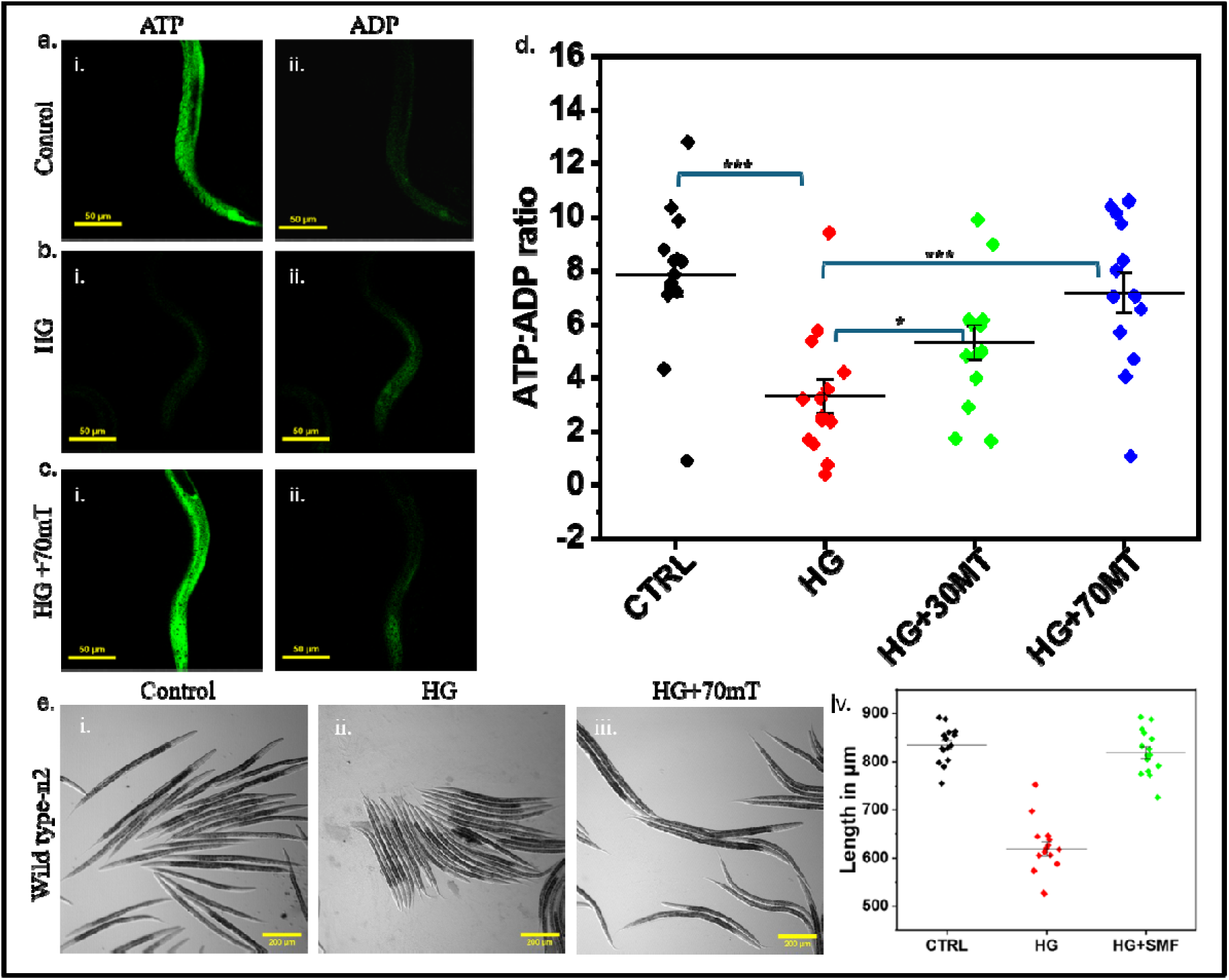
Effect of high glucose on C. elegans cellular energy environment and developmental biology. (a-i-ii) Shows ATP, ADP related fluorescence in the body of a live C. elegans in control condition**. (b-i-ii)** Shows ATP, ADP related fluorescence in the body of a live C. elegans after exposure to high glucose. **(c-i-ii)** Shows ATP and ADP levels in high glucose-exposed C. elegans after magnetic field treatment. **(d)** Quantitative comparison of ATP:ADP ratio in live C. elegans in different experimental conditions. **(e-i-iii)** C. elegans body size from control, high glucose exposed and magnetic field treated populations. **(e-iv)** Quantitative comparison of C. elegans body sizes in different experimental conditions.

### 4. Effect of low energy environment of fluorescence reporters of chaperones and immunity

Given the substantial reduction in cellular ATP levels observed under hyperglycaemic conditions, we next investigated whether stress-response signalling pathways involved in protein quality control were similarly affected. Molecular chaperones play a critical role in maintaining proteostasis by assisting protein folding and preventing the accumulation of damaged or misfolded proteins, particularly under conditions of oxidative stress [30]. However, the induction and activity of these pathways are energetically demanding processes that may be compromised in a low-ATP environment. To investigate this possibility, we employed the CL2070 strain expressing the hsp-16.2::GFP reporter. As shown in **(Figure 4a-i–iii),** hyperglycaemic worms exhibited a marked reduction in HSP-16.2 reporter fluorescence compared to control animals. This decrease may reflect impaired stress-response capacity resulting from the combined effects of elevated ROS and reduced cellular energy availability. Interestingly, magnetic field treatment restored reporter fluorescence to levels approaching those observed in control populations, consistent with the recovery of mitochondrial function, reduction of oxidative stress, and restoration of ATP levels described in previous sections. Quantitative analysis confirmed a significant reduction in HSP-16.2 reporter intensity following glucose exposure (p < 0.0001), which was subsequently rescued by magnetic field treatment **(Figure 4a-iv).** A similar trend was observed in worms expressing the t24b8.5 promoter reporter. T24b8.5 is a well known C. elegans gene that serves as a primary biomarker of the p38 Mitogen-Activated Protein Kinase (MAPK) innate immune pathway. Its proper activation can be coorelated with a robust immune response in response to external infections. Hyperglycaemic worms displayed significantly reduced t24b8.5 promoter reporter fluorescence compared to control populations (p < 0.0001) **(Figure 4b-i–iii),** suggesting suppression of stress-responsive signalling under conditions of metabolic dysfunction. T24b8.5 is directly downstream to transcription factor ATF-7 which functions as a key regulator of innate immune responses in C. elegans [31], and its activity is closely linked to cellular stress adaptation. To determine whether this reduced immune signalling during metabolic dysfunction extends upstream, we decided to check fluorescence from ATF-7::GFP based reporter. Consistent with the previous observations, hyperglycaemic animals exhibited a significant reduction in reporter fluorescence intensity from their nuclei compared to control worms (p < 0.0001) **(Figure 4c-i– iii).** Restoration of reporter expression was observed following magnetic field treatment, mirroring the recovery seen in both HSP-16.2 and T24b8.5 reporter strains. Collectively, these findings demonstrate that hyperglycaemia not only disrupts mitochondrial morphology, redox balance, lysosomal function, and cellular energetics, but also suppresses stress-response and immune-associated signalling pathways. Restoration of reporter activity following magnetic field treatment suggests that recovery occurs at multiple levels of biological organization, extending beyond organelle morphology to encompass functional cellular responses. Given the central role of ATP-dependent chaperones and stress-response pathways in maintaining proteostasis, we next investigated whether the hyperglycaemia-induced energetic deficit resulted in the accumulation of aggregation-prone proteins.

**Figure 4:**
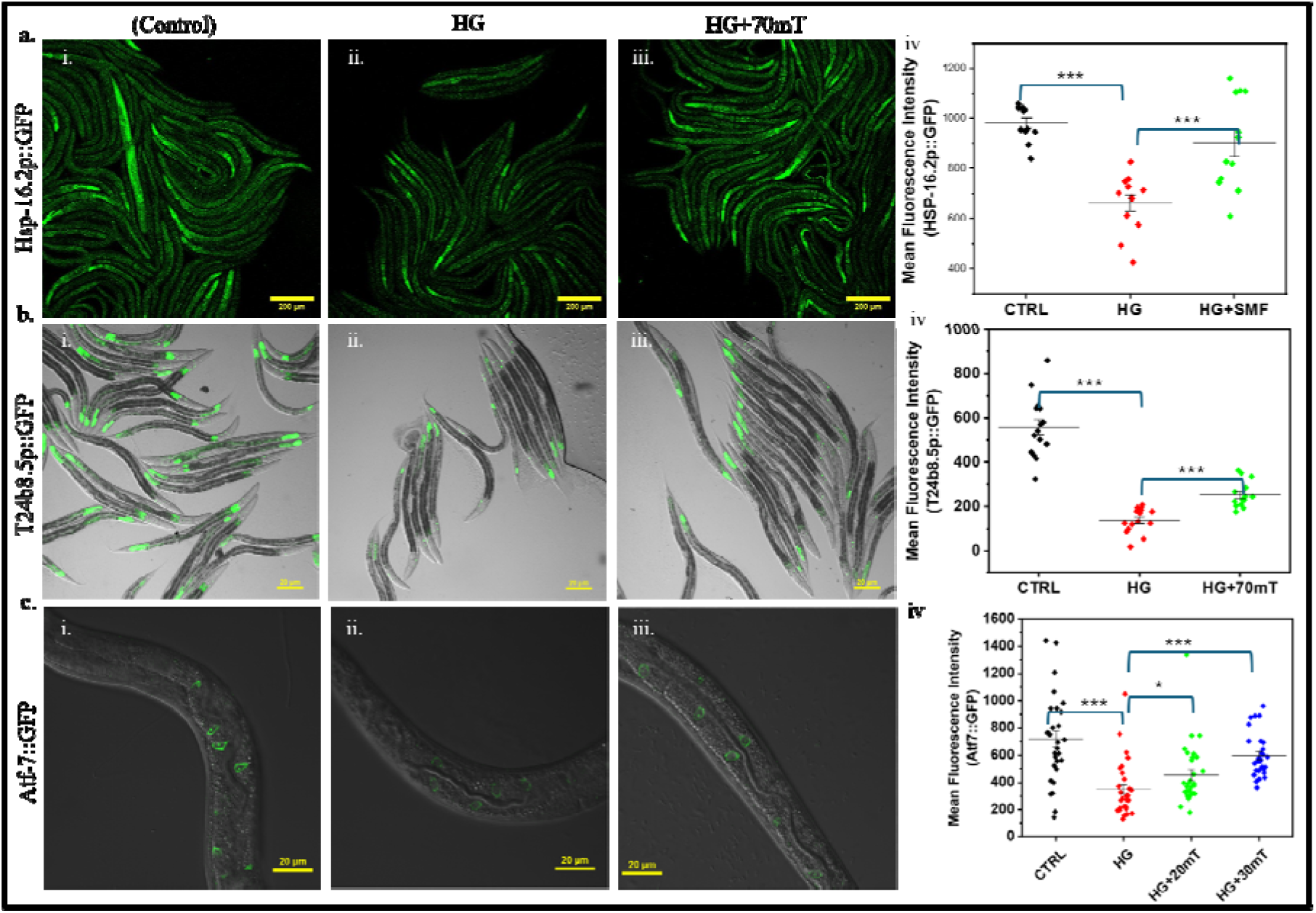
Effect of high glucose on C. elegans cellular energy environment and developmental biology. (a-i-iii) Shows hsp-16.2p::GFP based fluorescence in control, high glucose exposed and magnetic field treated worms previously exposed to high glucose **(a-iv)** Quantitative comparison of hsp-16.2p::GFP based fluorescence in C. elegans in different experimental conditions. **(b-i-iii)** Shows t24b8.5p based fluorescence in control, high glucose exposed and magnetic field treated worms previously exposed to high glucose. **(b-iv)** Quantitative comparison of t24b8.5p based fluorescence in C. elegans in different experimental conditions. **(c-i-iii)** Shows Atf-7::GFP based fluorescence in control, high glucose exposed and magnetic field treated worms previously exposed to high glucose **(c-iv)** Quantitative comparison of Atf-7::GFP based fluorescence in C. elegans in different experimental conditions.

### 5. Effect of impaired chaperone activity on proteotoxic clearance

Given the reduction in HSP-16.2 reporter fluorescence observed following glucose exposure, we next investigated whether the compromised stress-response environment translated into defects in protein quality control and proteostasis. Hyperglycaemia has previously been associated with increased protein aggregation and impaired clearance of damaged proteins, particularly under conditions of elevated oxidative stress and mitochondrial dysfunction. Therefore, in continuation with the phenomena observed in the previous sections, we examined the accumulation of aggregation-prone proteins within the body of C. elegans. For our first readout, we employed the EAK103 strain expressing polyglutamine-expanded Q138::YFP in body wall muscle cells. Polyglutamine proteins are highly prone to aggregation and are commonly used as reporters of proteostatic health. Consistent with the reduction in chaperone-associated reporter activity and the establishment of a low-energy, high-ROS environment, hyperglycaemic worms exhibited a marked increase in Q128::YFP aggregation compared to control populations (**Figure 5a-i–iii**). Large fluorescent aggregates were readily visible throughout the body wall musculature of glucose-exposed animals. Quantitative analysis revealed that the average number of large aggregates per worm increased substantially from approximately 3 in control animals to nearly 5 following glucose exposure. Notably, this phenotype was largely rescued by 70 mT magnetic field treatment, with aggregate numbers decreasing to slightly above 3, approaching control levels **(Figure 5a-iv).** To determine whether this effect extended to additional aggregation-prone proteins, we next examined worms expressing the soluble Q15::YFP reporter. Under normal physiological conditions, Q15 remains largely diffuse and does not typically form stable aggregates. However, hyperglycaemic worms exhibited increased accumulation and retention of Q15 fluorescence within muscle cells **(Figure 5b-i–iii),** suggesting a broader impairment in protein turnover and clearance mechanisms rather than simply enhanced aggregation of highly aggregation-prone proteins. Another strain DCD214 expressing tagRFP::pab-1 was used to assess the aggregation of non-neurodegenerative proteins. Pab-1 proteins naturally aggregate within the body of C. elegans with response to age or external cellular stress. In our study the RFP reporter would provide us with the fluorescence signal to show the extent of aggregation of pab-1. In our results we observed a significant increase pab-1 based RFP fluorescence in C. elegans exposed to 40mM hgh glucose compared to controlled worms (p=0.0005). the condition was rescued in populations exposed to 70mT magnetic field (p<0.0001). Taken together, these findings suggest that hyperglycaemia-induced mitochondrial dysfunction initiates a cascade of downstream events involving oxidative stress, ATP depletion, suppression of stress-response pathways, and ultimately impaired proteostasis. The accumulation of Q138, and even normally soluble Q15 proteins demonstrates that the consequences of metabolic dysfunction extend beyond organelle damage to affect global protein homeostasis. Restoration of protein clearance following magnetic field exposure further supports the central role of mitochondrial recovery in mediating the beneficial effects observed throughout this study. These results indicate that static magnetic fields not only repair mitochondrial morphology and restore cellular energetics but also alleviate downstream proteotoxic stress, highlighting their potential as a non-invasive strategy for combating aggregation-associated pathologies linked to metabolic and age-related disorders.

**Figure 5:**
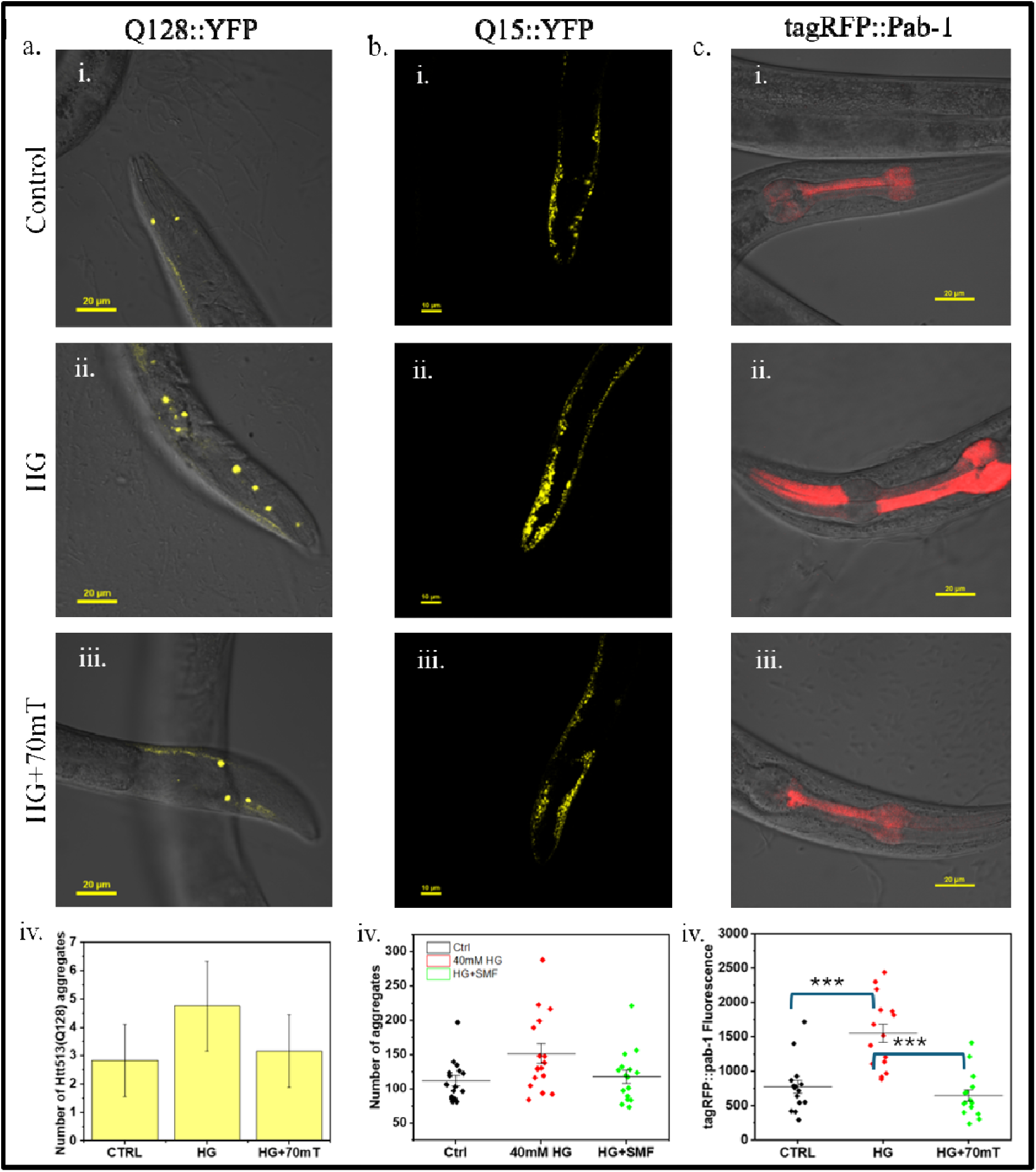
Effect of high glucose on C. elegans cellular energy environment and developmental biology. (a-i-iii) Shows aggregation of Q-128::YFP in control, high glucose exposed and magnetic field treated worms previously exposed to high glucose **(a-iv)** Quantitative comparison of number of aggregates of Q-128::YFP in C. elegans in different experimental conditions. **(b-i-iii)** Variation in expression of Q15::YFP in body wall m7uscle in C. elegans in control high glucose exposed and magnetic field treated populations previously exposed to high glucose. **(b-iv)** Quantitative comparison of difference in fluorescence based on Q-15::YFP expression in C. elegans in different experimental conditions. **(c-i-iii)** Aggregation of pab-1::RFP after high glucose treatment and its reversal in magnetic field exposed populations. **(c-iv)** Quantitative comparison of number of aggregates of pab-1::RFP in C. elegans in different experimental conditions.

## Discussion

The present study demonstrates that appropriately tuned static magnetic fields (SMFs) can reverse multiple downstream consequences of chronic hyperglycaemic stress in C. elegans, extending from mitochondrial dysfunction to organismal physiology. To model severe hyperglycaemia, worms were exposed to 40 mM glucose, a concentration previously shown to produce an internal glucose level of approximately 14 mM, comparable to severe hyperglycaemia in humans [27]. This relatively high glucose concentration was selected to reproduce, within the 2–3 day experimental timescale of C. elegans, the cumulative metabolic stress that develops over months or years in diabetic patients. Using SRRF and confocal microscopy together with functional reporter strains, we systematically evaluated mitochondrial morphology, oxidative stress, lysosomal integrity, cellular energetics, developmental progression, stress-response signalling, and proteostasis following exposure to different SMF strengths. Among the field strengths tested, 70 mT consistently produced the greatest recovery across all parameters, identifying an optimal therapeutic window. Hyperglycaemia enhances electron transport chain activity, resulting in excessive reactive oxygen species (ROS) production and mitochondrial fragmentation [7–12]. Consistent with this mechanism, hyperglycaemic worms displayed reduced mitochondrial area and perimeter together with increased circularity (Figure 1). SMF treatment progressively restored mitochondrial morphology between 20 and 70 mT, whereas 100 mT again induced fragmentation, indicating that the biological response is strongly field-strength dependent. Since maintenance of an interconnected mitochondrial network is essential for efficient oxidative phosphorylation and cellular homeostasis [27], restoration of mitochondrial morphology likely reflects improved mitochondrial function. Furthermore, under near-hypoxic stress induced by 60–70% Pluronic F-127, SMF-treated hyperglycaemic worms retained substantially larger mitochondrial structures than untreated animals, demonstrating enhanced mitochondrial resilience (Figure 1).

To our knowledge, this is the first in vivo demonstration of restoration of mitochondrial network morphology following hyperglycaemic damage by static magnetic fields.

Previous studies have reported that SMFs reduce intracellular ROS [28]. Consistent with these observations, 70 mT SMF significantly reduced gst-4p::GFP expression in hyperglycaemic worms while simultaneously normalizing lysosomal morphology and abundance (Figure 2). Because lysosomal dysfunction is closely linked to chronic oxidative stress and mitochondrial impairment [13], these findings suggest that restoration of mitochondrial homeostasis extends to downstream cellular quality-control pathways rather than acting solely through antioxidant effects. Restoration of mitochondrial morphology was accompanied by recovery of cellular bioenergetics. Although ATP depletion following hyperglycaemia has been demonstrated primarily in cultured mammalian cells [29], direct in vivo visualization of this energetic collapse has been lacking. Hyperglycaemic worms exhibited a marked reduction in ATP ratio despite excess glucose availability, whereas 70 mT SMF restored ATP levels almost to those of untreated controls (Figure 3). This bioenergetic recovery coincided with improved developmental progression and body size, establishing a functional link between mitochondrial structural recovery and organismal physiology. Chronic glucose exposure also suppressed the expression of hsp-16.2p::GFP together with the stress- and immune-responsive reporters atf-7 and T24B8.5 (Figure 4), consistent with impaired chaperone expression reported in diabetic models [30]. Restoration of these reporters following SMF treatment occurred concurrently with recovery of mitochondrial morphology, ATP production, and reduced oxidative stress, suggesting that improved cellular bioenergetics re-establishes protective stress signalling and cellular resilience. Loss of proteostasis is a major downstream consequence of mitochondrial dysfunction. Hyperglycaemic worms accumulated significantly higher levels of PAB-1 and Q138, aggregates, while even the normally soluble Q15 reporter formed abnormal inclusions (Figure 5), consistent with previous reports linking hyperglycaemia to protein aggregation [31, 32]. SMF treatment markedly reduced aggregate accumulation across all reporter strains, coinciding with restoration of mitochondrial morphology, ATP homeostasis, lysosomal integrity, and stress-response signalling, indicating coordinated recovery of cellular protein quality-control mechanisms. Previous magnetic field studies in diabetic models have primarily reported reductions in ROS and restoration of mitochondrial number using a 3 mT magnetic field combined with a 7 kV electric field [33], reversal of autophagic burden using 20–60 mT SMFs combined with intermittent fasting [34] or improved glycaemic control following continuous exposure to 100 mT SMFs [35]. In contrast, our study demonstrates, for the first time at the whole-organism level, that appropriately tuned SMFs restore mitochondrial network morphology and resilience following hyperglycaemic stress while simultaneously reversing downstream defects in oxidative stress, lysosomal homeostasis, ATP production, stress signalling, proteostasis, and organismal development. Collectively, these findings identify restoration of mitochondrial integrity as a central event underlying SMF-mediated recovery and establish 70 mT as the most effective field strength tested, highlighting the therapeutic potential of static magnetic fields as a non-invasive strategy for mitigating hyperglycaemia-induced cellular dysfunction.

## Conclusion

Hyperglycaemia-induced metabolic dysfunction is increasingly recognized as a major contributor to ageing-associated pathologies through its detrimental effects on mitochondrial homeostasis, oxidative stress regulation, cellular energetics, and proteostasis. In the present study, we demonstrate that exposure of Caenorhabditis elegans to high glucose results in widespread subcellular dysfunction characterized by mitochondrial fragmentation, elevated oxidative stress, lysosomal impairment, reduced ATP ratios, delayed development, suppression of stress-response signalling, and accumulation of aggregation-prone proteins. Importantly, static magnetic field (SMF) exposure produced a robust and dose-dependent reversal of these defects. Among the field strengths tested, 70 mT consistently provided the greatest benefit, restoring mitochondrial network architecture, improving mitochondrial resilience to secondary stress, reducing ROS accumulation, normalizing lysosomal morphology, and recovering cellular energy status. These improvements were accompanied by enhanced developmental progression, restoration of stress-responsive and immune-associated reporter activity, and a marked reduction in polyglutamine protein aggregation. In contrast, higher magnetic field strengths became detrimental, highlighting the importance of optimizing exposure parameters. Collectively, our findings support the concept that mitochondrial dysfunction represents a central driver of hyperglycaemia induced cellular pathology and that restoration of mitochondrial homeostasis can alleviate multiple downstream consequences of metabolic stress. The ability of static magnetic fields to improve mitochondrial function, cellular energetics, and proteostasis without pharmacological intervention highlights their potential as a non-invasive therapeutic strategy for metabolic disorders and age-associated neurodegenerative conditions. Further studies in mammalian systems will be required to establish the underlying mechanisms and evaluate translational applicability.

## Author Contributions

BM conceived and designed the experiments with inputs from CKN and LB, optimized experimental protocols, performed all *C. elegans* related experiments, imaging, and data analysis, and also wrote the manuscript. SC helped in conceiving the idea for the manuscript. AS helped in instrument handling during imaging. CKN and LB supervised the overall project, offered continuous guidance throughout the research, and contributed to the conceptual development and editing of the manuscript.

## Data Availability Statement

The data that support the findings of this study are available in the supplementary material of this article.

## Acknowledgements

The authors thank the Advanced Material Research Centre (AMRC), IIT Mandi and the Indian Knowledge System for Mental Health Application (IKSMHA), IIT Mandi for providing the facilities like cell culture, *C. elegans* model system and the sophisticated instruments necessary for the project. BM, SC, thanks the Ministry of Education, India (MoE), for the scholarship. AI tools were used for grammar correction, language refinement, and improving the clarity of the manuscript, also used to assist in developing the graphical concept and visual design of the Table of Contents (TOC) scheme.

## Supplementary Information

The Supporting Information mitochondrial morphology changes at different magnetic field strengths. Supplementary Figures S1-S4 provide additional evidence for the same

## References

1. Magliano, D.J., Boyko, E.J. and Atlas, D., 2025. 3. The global picture of diabetes. In Diabetes Atlas [Internet]*. 11th edition*. International Diabetes Federation.

2. Islam, K., Islam, R., Nguyen, I., Malik, H., Pirzadah, H., Shrestha, B., Lentz, I.B., Shekoohi, S. and Kaye, A.D., 2025. Diabetes mellitus and associated vascular disease: pathogenesis, complications, and evolving treatments. Advances in therapy, 42(6), pp.2659–2678.

3. Zhao, L., Yuan, J., Yang, Q., Ma, J., Yang, F., Zou, Y., Liu, K. and Liu, F., 2026. Diabetes and its complications: molecular mechanisms, prevention and treatment. Signal Transduction and Targeted Therapy, 11(1), p.22.

4. Szablewski, L., 2025. Associations between diabetes mellitus and neurodegenerative diseases. International journal of molecular sciences, 26(2), p.542.

5. Zhang, T., Yang, Y., Jiang, J., Du, W., Huang, G., Du, D. and Tao, S., 2025. The role of glucose metabolism in wound healing: an overview. Burns & Trauma, 13, p.tkaf053.

6. Chia, C.W., Egan, J.M. and Ferrucci, L., 2018. Age-related changes in glucose metabolism, hyperglycemia, and cardiovascular risk. Circulation research, 123(7), pp.886–904.

7. Rolo, A.P. and Palmeira, C.M., 2006. Diabetes and mitochondrial function: role of hyperglycemia and oxidative stress. Toxicology and applied pharmacology, 212(2), pp.167–178.

8. Halimi, S. and Vergès, B., 2014. Adverse effects and safety of SGLT-2 inhibitors. Diabetes & metabolism, 40(6), pp.S28–S34.

9. Poljšak, B., Kovač, V. and Milisav, I., 2022. Current uncertainties and future challenges regarding NAD+ boosting strategies. Antioxidants, 11(9), p.1637.

10. Sarangarajan, R.S.P.G., Meera, S., Rukkumani, R., Sankar, P. and Anuradha, G., 2017. Antioxidants: Friend or foe?. Asian Pacific journal of tropical medicine, 10(12), pp.1111–1116.

11. Sun, Z., Zhu, K., Zhao, W., Fei, X.F., Shi, L. and Zhang, Y., 2025. Potential mechanisms and clinical applications of static magnetic field therapy in glioma. Frontiers in Neurology, 16, p.1594874.

12. Zhang, C., Dong, C., Liu, X., Zhang, J., Li, Q., Chen, S., Zhao, H. and Huang, D., 2025. Recent Studies on the Effects of Static Magnetic Fields (SMF) on Reproductive Function. Current Issues in molecular biology, 47(2), p.116.

13. Beutner, G., Yuh, H.J., Goldenberg, I., Wallace, D.C., Porter Jr, G.A., Moss, A.J. and Sheu, S.S., 2025. Low magnetic fields stimulate cardiac mitochondrial bioenergetics with a bell-shaped response: Possibly via a radical pair mechanism. Computational and Structural Biotechnology Journal.

14. Tota, M., Jonderko, L., Witek, J., Novickij, V. and Kulbacka, J., 2024. Cellular and molecular effects of magnetic fields. International Journal of Molecular Sciences, 25(16), p.8973.

15. Saletnik, B., Saletnik, A., Słysz, E., Zaguła, G., Bajcar, M., Puchalska-Sarna, A. and Puchalski, C., 2022. The static magnetic field regulates the structure, biochemical activity, and gene expression of plants. Molecules, 27(18), p.5823.

16. Ren, J., Wang, M., Zhao, C., Luo, Y. and Tian, L., 2025. Mitochondria as key targets underlying hypomagnetic field-induced biological effects. iScience.

17. Carter, C.S., Huang, S.C., Searby, C.C., Cassaidy, B., Miller, M.J., Grzesik, W.J., Piorczynski, T.B., Pak, T.K., Walsh, S.A., Acevedo, M. and Zhang, Q., 2020. Exposure to static magnetic and electric fields treats type 2 diabetes. Cell metabolism, 32(4), pp.561–574.

18. Wang, Y., Feng, C., Yu, B., Wang, J., Chen, W., Song, C., Ji, X., Guo, R., Cheng, G., Chen, H. and Wang, X., 2024. Enhanced effects of intermittent fasting by magnetic fields in severe diabetes. Research, 7, p.0468.

19. Yu, B., Liu, J., Cheng, J., Zhang, L., Song, C., Tian, X., Fan, Y., Lv, Y. and Zhang, X., 2021. A static magnetic field improves iron metabolism and prevents high-fat-diet/streptozocin-induced diabetes. The Innovation, 2(1).

20. Yuan, Q., Zeng, Z.L., Yang, S., Li, A., Zu, X. and Liu, J., 2022. Mitochondrial stress in metabolic inflammation: modest benefits and full losses. Oxidative Medicine and Cellular Longevity, 2022(1), p.8803404.

21. Rizwan, H., Pal, S., Sabnam, S. and Pal, A., 2020. High glucose augments ROS generation regulates mitochondrial dysfunction and apoptosis via stress signalling cascades in keratinocytes. Life sciences, 241, p.117148.

22. Yu, T., Jhun, B.S. and Yoon, Y., 2011. High-glucose stimulation increases reactive oxygen species production through the calcium and mitogen-activated protein kinase-mediated activation of mitochondrial fission. Antioxidants & redox signaling, 14(3), pp.425–437.

23. Zong, Y., Li, H., Liao, P., Chen, L., Pan, Y., Zheng, Y., Zhang, C., Liu, D., Zheng, M. and Gao, J., 2024. Mitochondrial dysfunction: mechanisms and advances in therapy. Signal transduction and targeted therapy, 9(1), p.124.

24. Liu, S., Liu, J., Wang, Y., Deng, F. and Deng, Z., 2025. Oxidative stress: signaling pathways, biological functions, and disease. MedComm, 6(7), p.e70268.

25. De la Mata, M., Cotán, D., Villanueva-Paz, M., De Lavera, I., Álvarez-Córdoba, M., Luzón-Hidalgo, R., Suárez-Rivero, J.M., Tiscornia, G. and Oropesa-Ávila, M., 2016. Mitochondrial dysfunction in lysosomal storage disorders. Diseases, 4(4), p.31.

26. Plotegher, N. and Duchen, M.R., 2017. Mitochondrial dysfunction and neurodegeneration in lysosomal storage disorders. Trends in molecular medicine, 23(2), pp.116–134.

27. Ludlaim, A.M., Waddington, S.N. and McKay, T.R., 2025. Unifying biology of neurodegeneration in lysosomal storage diseases. Journal of Inherited Metabolic Disease, 48(1), p.e12833.

28. Frachini, E.C.G., Silva, J.B., Fornaciari, B., Baptista, M.S., Ulrich, H. and Petri, D.F.S., 2024. Static magnetic field reduces intracellular ROS levels and protects cells against peroxide-induced damage: suggested roles for catalase. Neurotoxicity Research, 42(1), p.2.

29. Knudsen, J.G., Hamilton, A., Ramracheya, R., Tarasov, A.I., Brereton, M., Haythorne, E., Chibalina, M.V., Spegel, P., Mulder, H., Zhang, Q. and Ashcroft, F.M., 2019. Dysregulation of glucagon secretion by hyperglycemia-induced sodium-dependent reduction of ATP production. Cell metabolism, 29(2), pp.430–442.

30. de Oliveira, A.A., Mendoza, V.O., Rastogi, S. and Nunes, K.P., 2022. New insights into the role and therapeutic potential of HSP70 in diabetes. Pharmacological research, 178, p.106173.

31. Lv, Y.Q., Yuan, L., Sun, Y., Dou, H.W., Su, J.H., Hou, Z.P., Li, J.Y. and Li, W., 2022. Long-term hyperglycemia aggravates α-synuclein aggregation and dopaminergic neuronal loss in a Parkinson’s disease mouse model. Translational Neurodegeneration, 11(1), p.14.

32. Talaei, F., Van Praag, V.M., Shishavan, M.H., Landheer, S.W., Buikema, H. and Henning, R.H., 2014. Increased protein aggregation in Zucker diabetic fatty rat brain: identification of key mechanistic targets and the therapeutic application of hydrogen sulfide. BMC cell biology, 15(1), p.1.

33. Carter, C.S., Huang, S.C., Searby, C.C., Cassaidy, B., Miller, M.J., Grzesik, W.J., Piorczynski, T.B., Pak, T.K., Walsh, S.A., Acevedo, M. and Zhang, Q., 2020. Exposure to static magnetic and electric fields treats type 2 diabetes. Cell metabolism, 32(4), pp.561–574.

34. Wang, Y., Feng, C., Yu, B., Wang, J., Chen, W., Song, C., Ji, X., Guo, R., Cheng, G., Chen, H. and Wang, X., 2024. Enhanced effects of intermittent fasting by magnetic fields in severe diabetes. Research, 7, p.0468.

35. Yu, B., Liu, J., Cheng, J., Zhang, L., Song, C., Tian, X., Fan, Y., Lv, Y. and Zhang, X., 2021. A static magnetic field improves iron metabolism and prevents high-fat-diet/streptozocin-induced diabetes. The Innovation, 2(1).

