## Supplementary Information for "Magnetic Fields Improves the Cellular function in Hyperglycaemia"

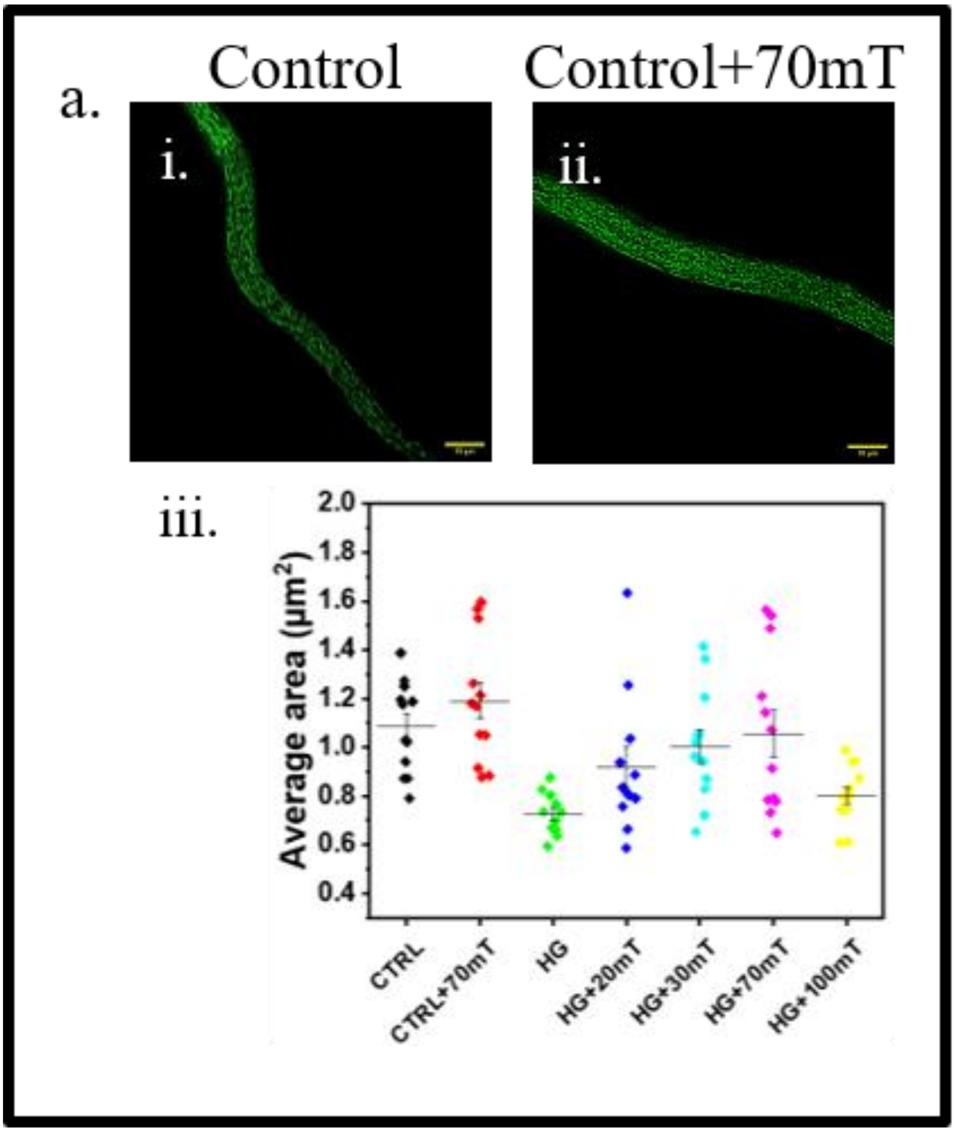


Figure S1: Effect of magnetic field on C. elegans mitochondria. (a-i-ii) Shows magnetic field induced improvement in mitochondrial morphology (iii) Quantitative analysis of improvement in overall mitochondrial area after application of magnetic field even in control specimens.

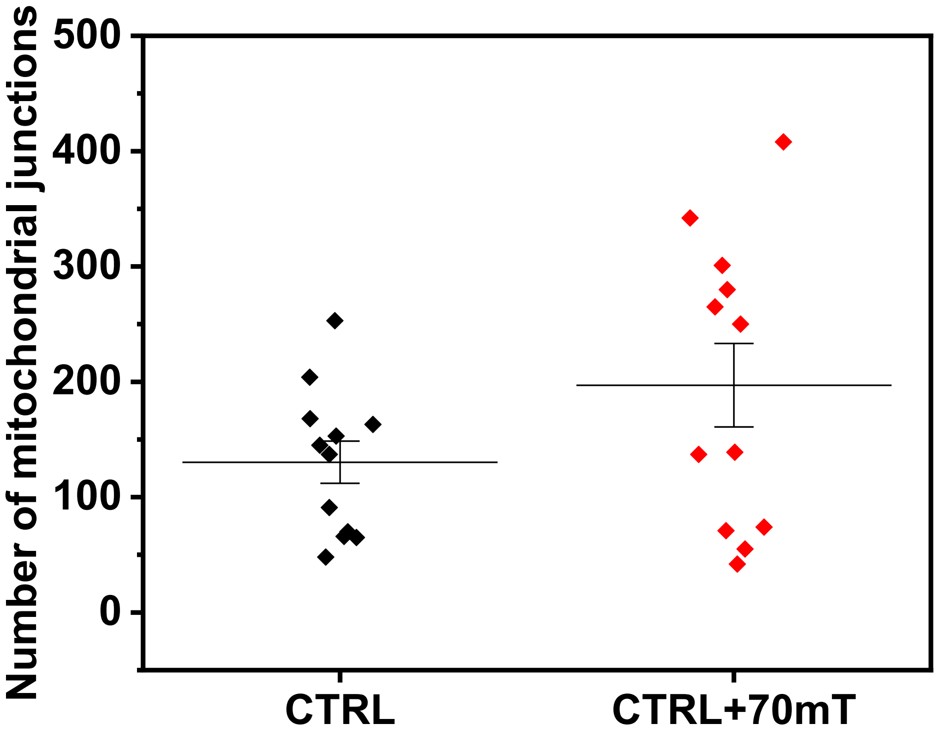


Figure S2: Increase in number of mitochondrial junctions after magnetic field treatment in control C. elegans

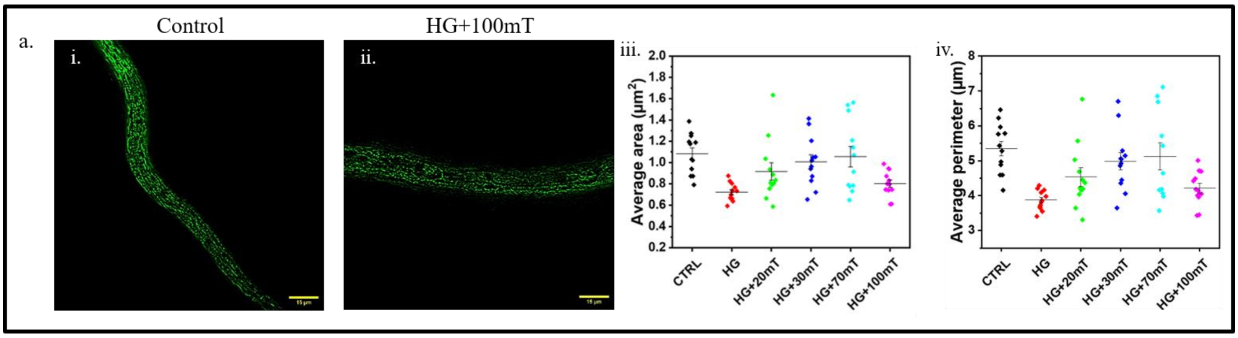


Figure S3: Disrupted mitochondrial health after a certain field strength (a-i-ii) Shows magnetic field induced disruption in mitochondrial morphology in high glucose exposed specimens (a-iii-iv) Average mitochondrial area (iii) and perimeter (iv) had reduced significantly in the 100 mT exposed C. elegans specimens with values Nealy equal to the high glucose exposed worms

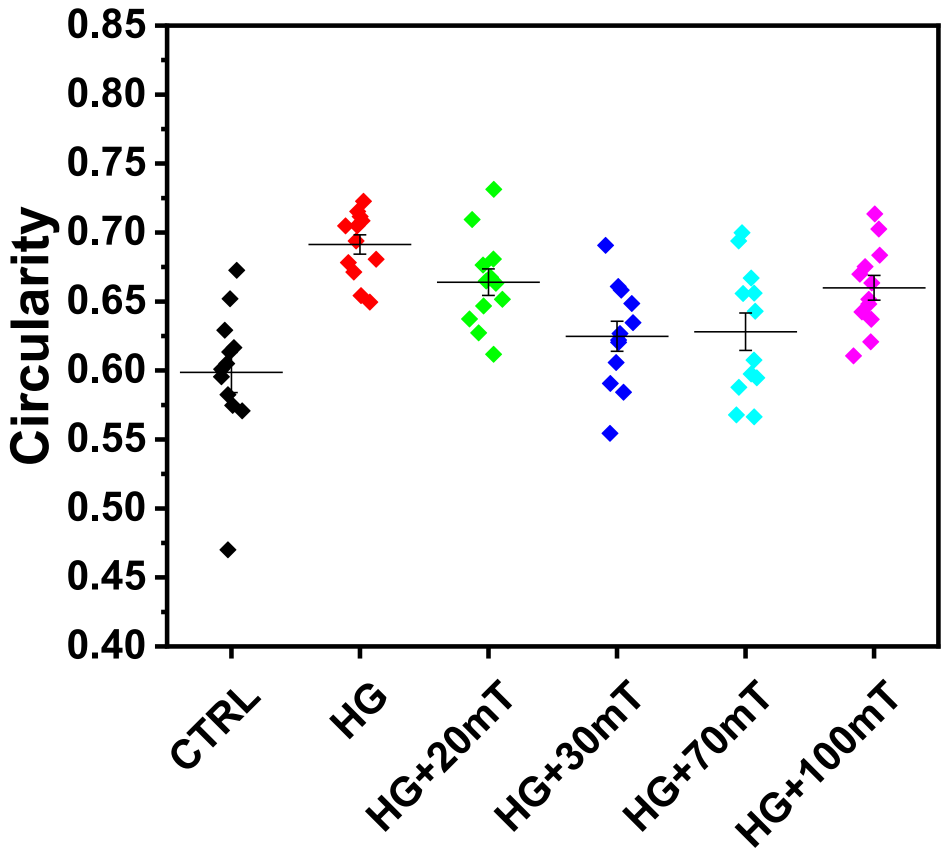


Figure S4: reduction in mitochondrial circularity in high glucose exposed C. elegans after treatment with SMF
